# Microglia influence neurofilament deposition in ALS iPSC-derived motor neurons

**DOI:** 10.1101/2021.12.30.474573

**Authors:** Reilly L. Allison, Jacob W. Adelman, Jenica Abrudan, Raul A. Urrutia, Michael T. Zimmermann, Angela J. Mathison, Allison D. Ebert

**Author notes:** Corresponding author: Allison Ebert, 8701 Watertown Plank Rd, Milwaukee, WI 53226.

## Abstract

Amyotrophic lateral sclerosis (ALS) is a fatal neurodegenerative disease in which upper and lower motor neuron loss is the primary phenotype, leading to muscle weakness and wasting, respiratory failure, and death. Although a portion of ALS cases are linked to one of over 50 unique genes, the vast majority of cases are sporadic in nature. However, the mechanisms underlying the motor neuron loss in either familial or sporadic ALS are not entirely clear. Here we used induced pluripotent stem cells derived from a set of identical twin brothers discordant for ALS to assess the role of astrocytes and microglia on the expression and accumulation of neurofilament proteins in motor neurons. We found that motor neurons derived from the affected twin exhibited increased transcript levels of all three neurofilament isoforms and increased expression of phosphorylated neurofilament puncta. We further found that treatment of the motor neurons with astrocyte conditioned medium and microglial conditioned medium significantly impacted neurofilament deposition. Together, these data suggest that glial-secreted factors can alter neurofilament pathology in ALS iPSC-derived motor neurons.

## Introduction

Amyotrophic lateral sclerosis (ALS) is a severe and always fatal neurodegenerative disease in which upper and lower motor neurons are lost leading to muscle weakness, paralysis, and death typically within 2-5 years of diagnosis. ALS has been causally linked to several different genetic mutations, but the vast majority of cases are still sporadic in nature with no known genetic cause [1]. Although various mechanisms have been postulated as leading to the motor neuron loss [1], there is still a lack of clear understanding of motor neuron degeneration in ALS.

Despite an unknown cause of motor neuron loss, intracellular inclusions of aggregated proteins are clear pathological hallmarks that span both sporadic and familial ALS patient samples [2]. Aggregates containing TDP-43 are found in greater than 90% of patients [3], but SOD1, optineurin (OPTN), ubiquitin, and neurofilament are also commonly found [2, 4, 5]. Protein aggregation may contribute to motor neuron death through activation of oxidative stress and/or by causing general cellular dysfunction through sequestering cytosolic proteins to the aggregates themselves [6-9]. The alteration of neurofilament level is of particular interest in ALS because neurofilament is found in high abundance in the cerebral spinal fluid from ALS patients and may serve as a key biomarker to ALS disease onset, progression, and therapeutic success [10-12].

Neurofilaments (NFs) are intermediate filament proteins consisting of heteropolymers of the heavy (H), medium (M), and light (L) subunits [13, 14]. NFs are abundantly expressed in neurons and they provide structural support, facilitate organelle transport, participate in intracellular signaling, and regulate axon growth [15, 16]. Mutations in *NEFH* have been found in a small number of ALS cases [5], but the role of NF proteins in ALS have more recently focused on the potentially pathogenic nature of NF neuronal accumulations [11]. For example, high expression of human NF-H proteins in mice caused large inclusions in motor neurons and contributed to axonal atrophy, diminished conductance, and severe motor deficits [17, 18]. Additionally, NF aggregates have been observed in human stem cell models of ALS from both sporadic and familial forms of the disease [19-21]. The mechanisms underlying NF aggregation in ALS are still not fully elucidated, but evidence suggests that excitotoxicity increases NF hyperphosphorylation in neurons [22] thereby promoting NF aggregation.

Increased neuronal excitotoxicity has long been a proposed mechanism associated with motor neuron loss in ALS [23, 24]. Astrocytes play a key role in reducing excess glutamate in the synaptic cleft that can lead to neuronal excitotoxicity, but evidence indicates that ALS astrocytes are dysfunctional and actively induce motor neuron toxicity [23, 24]. For example, the presence of ALS astrocytes and/or their secreted proteins are sufficient to induce motor neuron damage and loss both *in vitro* and *in vivo* through non-cell autonomous mechanisms [25-30]. Similarly, ALS microglia have been shown to exhibit altered morphology and functionality *in vitro* and *in vivo* [31-34]. Interestingly, malfunctioning microglia are also able to induce astrocyte toxicity [35] suggesting a tight interplay between glial cells in health and disease.

Considering the mostly sporadic nature of ALS, having appropriate model systems can be a challenge. While induced pluripotent stem cells (iPSCs) derived from sporadic ALS patients offer a unique opportunity to assess disease properties in specific patient backgrounds, they also prevent the use of isogenic controls as there are no specific gene mutations to correct. To circumvent this problem, we have previously generated and characterized iPSCs from identical twin brothers who are discordant for ALS [36]. In our previous studies, we did not find disease-relevant genetic variations between the brothers [36] suggesting that mechanisms at the cellular level may be more influential to disease processes. Therefore, here we used RNA sequencing on the iPSC-derived motor neurons to assess whether there were specific molecular pathways altered in the affected twin that could contribute to the affected brother’s ALS diagnosis. Although there were several transcriptional differences between the brothers, we focused our attention on neurofilament due to its relationship to ALS disease pathology. Interestingly, we found that the affected twin exhibited increased transcript expression of all three neurofilament subtypes as well as neurofilament aggregation in the iPSC-derived motor neurons. Moreover, we found that treating the affected motor neurons with microglia derived from the healthy brother’s iPSCs significantly reduced the neurofilament deposition. Together, these data suggest that glial-motor neuron interactions are important modulators of neurofilament pathology in ALS.

## Materials and Methods

### Pluripotent Stem Cells

iPSC lines were previously generated from identical twins discordant for ALS (HB19.2, AB34.12) under an approved IRB protocol (PRO00024167); details about the individuals, iPSC generation, and pluripotency characterization is described in our previous work [36]. iPSCs were maintained on Matrigel (Corning) in Essential 8 (Gibco) and passaged every 4-6 days. The iPSCs and differentiated cells were confirmed mycoplasma negative.

### Motor neuron differentiation and treatments

Spinal motor neurons were differentiated using a previously published protocol [37]. Briefly, embryoid bodies were generated from iPSCs and patterned in the presence of Chir-99021 with dual SMAD inhibition (SB 431542 and LDN 1931899) followed by treatment with retinoic acid (RA), smoothened agonist (SAG), and DAPT. Spinal motor neuron progenitor cells were then dissociated and plated on Matrigel-coated glass coverslips (ICC) or 6-well plates (protein/ RNA collection) for terminal differentiation and maturation in growth factor supplemented medium for 28-42 days *in vitro*. Treatments with ACM were performed with 20% ACM in MN Maturation media after day 28 and left for 48 hours at 37C before fixing, collection, or analyses.

### Glial differentiation and treatments

Spinal cord patterned astrocytes were generated from iPSC-derived neural progenitor cells (NPCs) [38]. Briefly, iPSCs were grown to confluency, dissociated with Accutase, and plated at 2 million cells/well into Matrigel (Corning) coated 6 well-plates in NPC base medium (50% DMEM/ F12, 50% Neurobasal, 2% B27, 1% N2, 1% Antibiotic/ Antimycotic, 0.1% B-mercaptoethanol, 50ng/mL Laminin, and 0.5uM Ascorbic Acid) supplemented with 10uM Y-27632, 3uM Chir-99021, 40uM SB431542, and 0.2uM LDN193189. On day 1, NPC media was changed and supplemented with 3uM Chir-99021, 40uM SB431542, and 0.2uM LDN193189. On Days 2 and 3, NPC media was supplemented with 3uM Chir-99021, 40uM SB431542, 0.2uM LDN193189, 100nM retinoic acid (RA), and 500nM hedgehog smoothened agonist (SAG). From day 4 onwards, NPC media was supplemented with 40uM SB431542, 0.2uM LDN193189, 100nM RA, and 500nM SAG and changed daily. NPCs were passaged via Accutase treatment on Day 6 (P1), Day 12 (P2), and Day 18 (P3). P3 NPCs were used for astrocyte differentiations and cultured in ScienCell Astrocyte Medium containing 1% Astrocyte Growth Supplement, 1% Penicillin/Streptomycin, and 2% B27. Cells were fed every 48 hours and passaged with Accutase every 6-9 days upon confluency (minimum of 3 passages). Passage 4 cells were considered fully differentiated and seeded onto T75 tissue culture flasks for ACM generation and collection.

Microglia were differentiated using the commerically available differentiation kit (STEMCELL Technologies #05310, #100-0019, #100-0020) based on a previously published protocol [39]. Briefly, iPSCs were differentiated into hematopoietic progenitor cells (HPCs) using the STEMdiff Hematopoietic Kit (StemCell Technologies). Floating HPCs were then collected and plated at 50,000 cells/mL in STEMdiff Microglia Differentiation media (StemCell Technologies) for 24 days, followed by rapid maturation in STEMdiff Microglia Maturation media (StemCell Technologies) for at least 4 days. MCM was collected after day 28, spun to remove cells or debris, and stored in sterile Falcon tubes at –20C. Frozen medias were slowly thawed on ice before use.

Astrocyte conditioned media was collected upon each media change after P4, spun to remove any cells or debris, and stored in sterile Falcon tubes at –20C. Frozen medias were slowly thawed on ice before use. Treatments with MCM were performed with 20% MCM in supplemented Astrocyte media for 48 hours before removal of treatment media. Cells were then rinsed with PBS and fed with supplemented Astrocyte media. Media was collected from treated astrocytes after 48 hours.

### RNA sequencing

Cell pellets were harvested at 3 weeks of differentiation from both affected (quadruplicate) and unaffected (duplicate) differentiations, and transcriptome (RNAseq) analysis was completed at MCW’s Genomic Science and Precision Medicine Center (GSPMC). Total RNA was isolated according to manufacturer’s instructions (Qiagen, RNeasy mini kit) with on-column DNA digestion and quality assessed by fragment analysis (Agilent). All samples were good quality RNA (RIN>8.3), and 500ng input was aliquoted for library preparation. Libraries were prepared according to manufacturer’s protocols (TruSeq stranded mRNA, Illumina) with a final quality check completed by fragment analysis (Agilent) and quantification by qPCR (Kapa Library Quantification Kit, Kapa Biosystems). Samples were multiplexed and sequenced on a NovaSeq6000 SP flow cell, with 2×100bp read lengths captured. Sequencing reads were processed through the MAPR-Seq Workflow [40] with differential expression analysis completed with Bioconductor, edgeR v 3.8.6 software [41]. Genes with a false discovery rate (FDR) less than 5% and an absolute fold change ≥ 2 were considered significantly differentially expressed. Data have been deposited in the Gene Expression Omnibus (GEO) database (GEO accession number GSE192755).

### Western Blot

Cell pellets were lysed by snap-freezing and vortexing with Triton X-100, then protein concentration was determined using a BCA assay (ThermoFisher). Equal amounts of protein were loaded onto 10% or 12% pre-cast Tris-HCl Mini-PROTEAN gels (Bio-Rad) and proteins separated by electrophoresis, then transferred to PVDF membranes (Bio-Rad). Membranes were blocked for 1 hour in Odyssey TBS Blocking Buffer (LI-COR) followed by overnight primary antibody incubation and 30 minute secondary antibody incubation. Quantification was performed with FIJI (ImageJ) and normalized to REVERT total protein stain (LI-COR). Primary antibodies used were rabbit anti-NF200 (Sigma, N4142, used at 1:2000 dilution), mouse anti-NF145 (Sigma, MAB1621, used at 1:500 dilution), and mouse anti-NF68 (Sigma, N5139, used at 1:500 dilution). Secondary antibodies used were anti-rabbit IRDye 800CW (LI-COR, 1:5000 dilution) and anti-mouse IRDye 680RD (LI-COR, 1:5000 dilution).

### Immunocytochemistry

Plated cells were fixed in 4% paraformaldehyde (PFA) for 20 minutes at room temperature and rinsed with PBS. Nonspecific labeling was blocked and the cells permeabilized with 0.25% Triton X-100 in PBS with 1% BSA and 0.1% Tween 20 for 15 minutes at room temperature. Cells were incubated with primary antibodies overnight at 4C, then labeled with appropriate fluorescently-tagged secondary antibodies. Hoechst nuclear dye was used to label nuclei. Primary antibodies used were goat anti-choline acetyltransferase (ChAT; Sigma, AB144P, 1:50 dilution), rabbit anti-neurofilament 200 (NF200; Sigma, N4142, 1:500 dilution), and mouse anti-phosphoNF (Sigma, MAB1592, 1:300 dilution). Secondary antibodies used were donkey anti-mouse AF488 (Invitrogen), goat anti-chicken AF568 (Invitrogen), donkey anti-rabbit AF546 (Invitrogen), and donkey anti-mouse AF647 (Invitrogen). All secondaries were used at 1:1000 dilution.

Analyses were performed on three randomly selected fields per coverslip using standard fluorescent microscopy and equivalent exposure conditions at 20x. Images were analyzed for total fluorescence in each channel using FIJI (ImageJ) software. NF200 and total phNF stains were normalized to ChAT in order to account for differences in cell densities between regions of interest. phNF+ aggregates were confirmed to colocalize with ChAT stain and were analyzed using Nikon Elements Object Count function. On the software, thresholds were set to identify aggregates as 3x greater intensity than average phNF stain intensity, and restrictions were set for size and circularity (area max=259.51um2, circularity min=0.50). phNF+ aggregates were represented as a percent of normalized phNF stain for each image to account for differences in cell densities between regions of interest. Representative images were acquired using a confocal microscope with a 63x oil objective and are displayed as the maximum intensity projection (MIP) of a z-stack of images.

### Multiplex cytokine array

Eve Technologies (Calgary, Alberta, Canada) performed the 48 multiplex cytokine array assay from duplicate differentiations using conditioned medium samples generated from iPSC-derived astrocytes and microglia.

### Statistical analyses

WB and ICC experiments were performed in triplicate on a minimum of two biological replicates. The multiplex cytokine array was performed on samples from two biological replicates for each condition. Data were analyzed using GraphPad Prism software and the appropriate statistical tests including the Student’s t-test and 1-way ANOVA followed by Tukey’s post hoc analysis of significance. Changes were considered statistically significant when p<0.05.

## Results

In order to model sporadic ALS, we have previously generated iPSCs from identical twin brothers discordant for ALS [36], which offers as close to an isogenic iPSC pair as possible. Our previous study used whole genome sequencing and found no substantial genetic differences or mutations in known ALS-associated genes between the brothers that would specifically indicate a genetic basis for the affected brother’s ALS diagnosis [36]. Moreover, we found no differences in motor neuron survival or response to glutamate stress when comparing iPSC-derived motor neurons between the affected and unaffected twins [36]. Therefore, here we used RNA sequencing on the motor neurons to get a better global picture of transcriptional signatures between the affected and unaffected cells to identify molecular pathways that may be involved in motor neuron malfunction and loss. We found 40,418 transcripts to be differentially expressed between the cells, with 3,086 significantly upregulated and 2,321 significantly downregulated in the affected motor neurons compared to the healthy control (Fig 1A). Pathway analysis identified Axonal Guidance Signaling and Synaptogenesis Signaling as the two most altered pathways with a -log(p-value) of 26.8 and 22.3, respectively. ALS Signaling, with a -log(p-value) of 11.2, was also highly altered based on pathway analysis. When assessing the genes associated with the ALS Signaling pathwy, we found that NF-H (NEFH), NF-M (NEFM), and NF-L (NEFL) were all significantly upregulated in the affected brother (AB) motor neurons compared to the healthy brother (HB) (Fig 1A, B); NF-H had a log2 fold change of 1.58 (p-adj = 0.0006), NF-M had a log2 fold change of 1.9 (p-adj = 0.0031), and NF-L had a log2 fold change of 1.81 (p-adj = 0.0006). In contrast, peripherin (PRPH), a neuronal intermediate filament protein found in neurons of the peripheral nervous system and in central nervous system neurons with projections to the periphery, was unchanged between the affected and unaffected motor neurons (Fig. 1A). Downstream analysis using Ingenuity Pathway Analysis (IPA) placed these altered NF transcripts in the ALS signaling pathway showing that upregulation of NFs can lead to neuronal protein inclusions, excitotoxicity, and motor neuron death (Fig. 1C).

**Figure 1:**
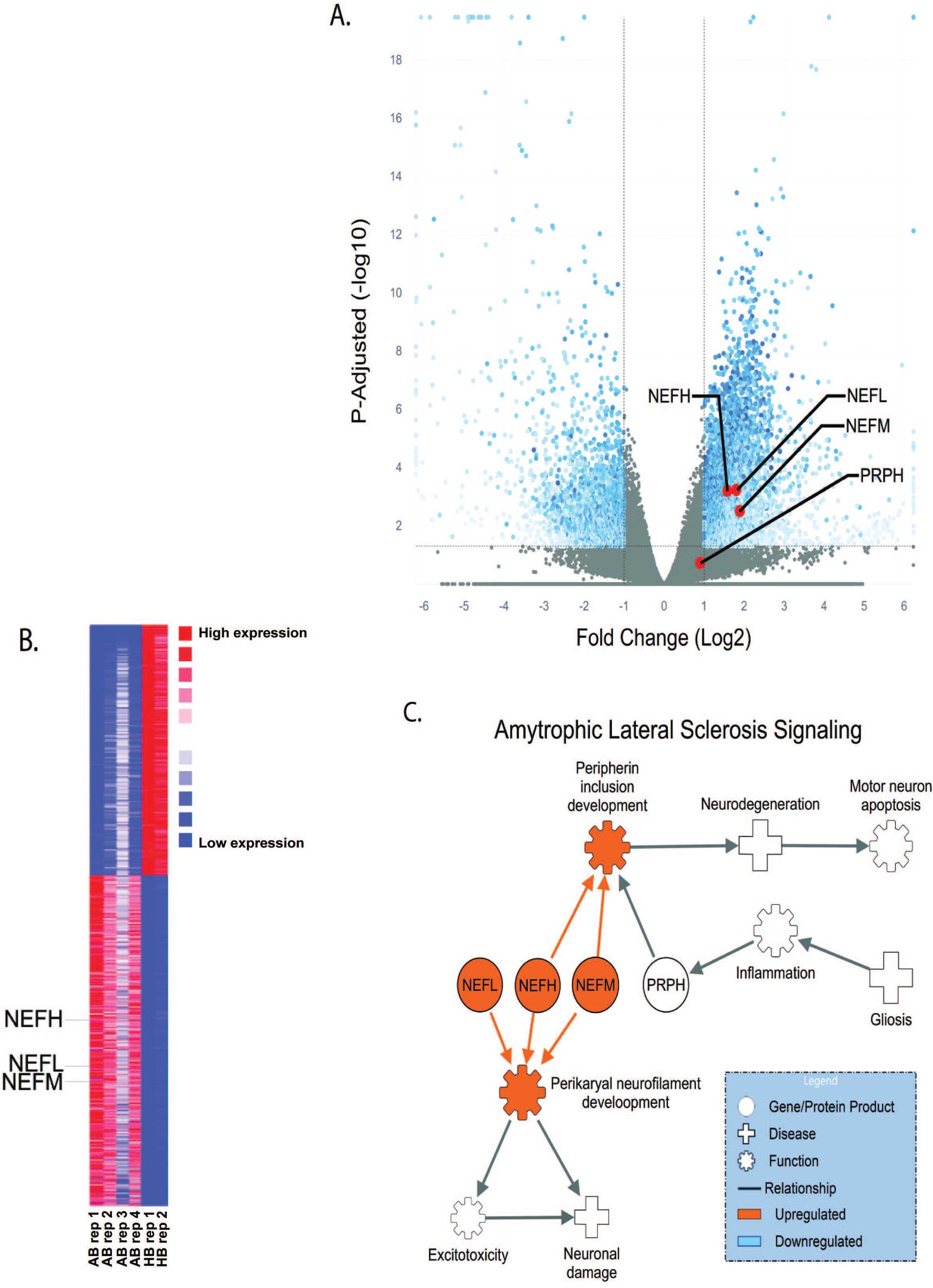
Comparing neurofilament transcripts and ALS-associated signaling in ALS-discordant twins. **(A)** Volcano plot and **(B)** heatmap of differential expression RNA sequencing (DESeq2) comparing the motor neuron transcriptome of an ALS-affected patient (AB) to his non-affected twin (HB). Neurofilament transcripts for neurofilamanet heavy (NEFH), neurofilament medium (NEFM), and neurofilament light (NEFL) are denoted with red dots and labeled [P_adj_ < 0.0501 FC ≥ ±1]. For comparison, peripherin (PRPH) was unchanged between the AB and HB. **(C)** Ingenuity pathway analysis (IPA) highlighting the role of altered neurofilament gene transcripts in ALS signaling [p < 0.01; FC > ±1.35].

With the observed increase in NF noted at the mRNA level, we then assessed protein levels of NF-H (NF200), NF-M (NF145), and NF-L (NF68) using both western blot and immunocytochemistry. Western blot analysis did not reveal a difference in NF protein levels in motor neurons between the AB and the HB for any of the NF subunits (Fig. 2A-D). Using immunocytochemistry we observed large accumulations of NF-200 staining along the neurites in the AB motor neuron cultures that were generally absent in the HB motor neurons (Fig. 2E, arrowheads); these data are similar to what has been observed in mutant SOD1 iPSC-derived motor neurons [20]. However, there was no difference in total NF200 (NF-H) expression in motor neurons between the AB and HB (Fig. 2F). We then used immunocytochemistry for phosphorylated NF (phNF) and found that although total phNF immunofluorescence was consistent between motor neurons from the AB and the HB (Fig. S1), there was a significant increase in the number of phNF positive puncta along the neurites and in the cell bodies of the AB iPSC-derived motor neurons compared to the HB motor neurons (Fig. 2E, G, arrows). These data indicate that NF accumulation may be a contributing factor to motor neuron demise in the AB.

**Figure 2:**
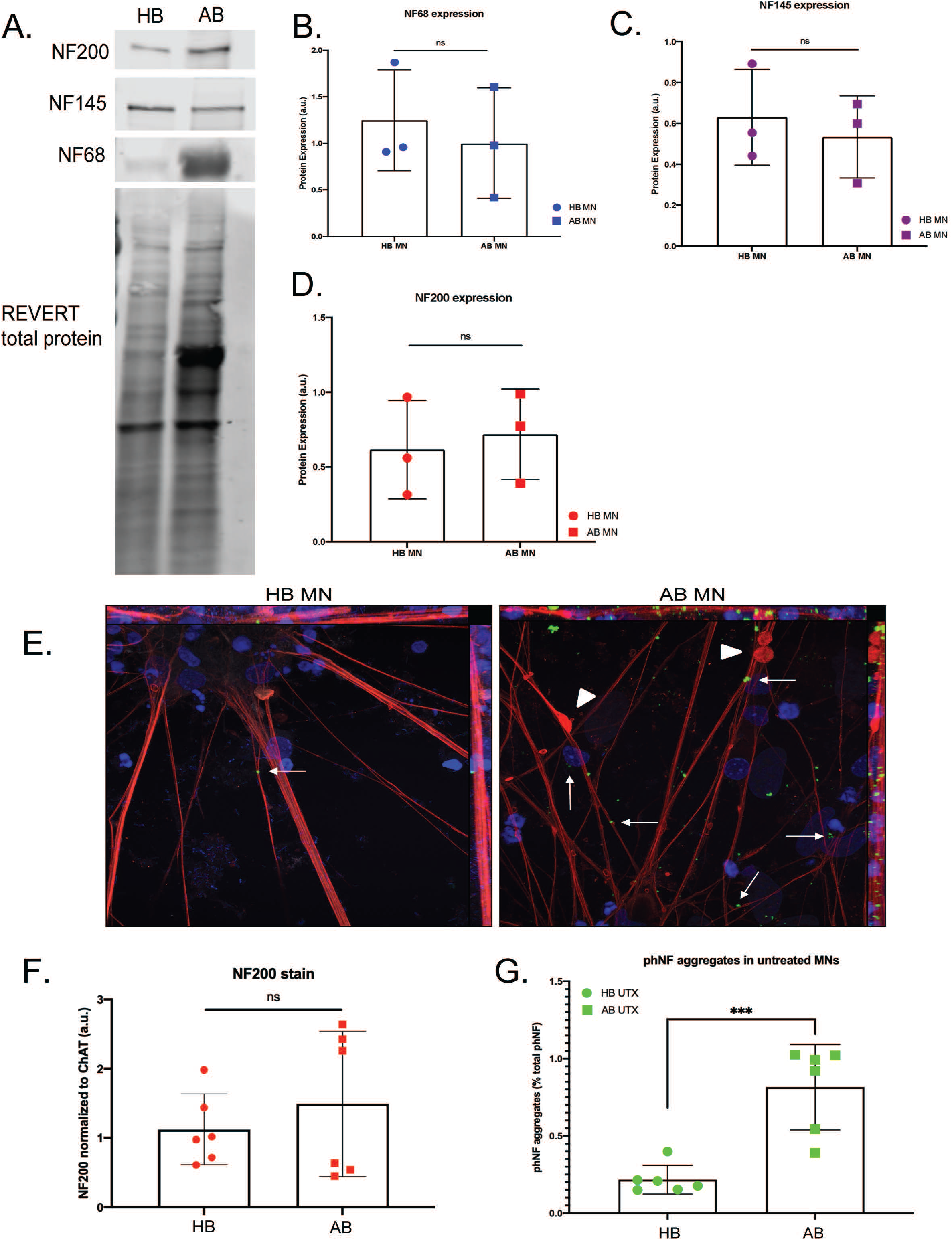
Aggregation of phosphorylated neurofilaments identified in AB motor neurons. (**A-D**) Western Blot (WB) for NF68 (NF-L) (**B**), NF145 (NF-M) (**C**), and NF200 (NF-H) (**D**) subunits reveal no significant differences between HB and AB MN protein expression. (**E**) Representative immunocytochemistry (ICC) images of HB (left) and AB (right) motor neurons (MNs) (red = NF200, green = phosphorylated NF (phNF), blue = DAPI; 63x objective). Arrows indicate aggregated phNF in neurites and cell bodies; arrowheads indicate neurite swellings. (**F**) Despite no significant differences between HB and AB for total NF200 expression, (**G**) quantification reveals significantly increased number of phNF+ aggregates in AB MNs compared to HB MNs [t-test, ***p<0.0005].

Considering that glial cells have been shown to contribute to ALS motor neuron toxicity and NF pathology (Fig. 1C), we next tested whether the presence of astrocyte and/or microglia conditioned medium (ACM and MCM, respectively) could impact NF expression or accumulation. We differentiated HB and AB iPSCs into astrocytes and microglia (Fig. S2) and analyzed the ACM and MCM with a multiplex cytokine array (Fig. 3). We then focused our analysis on the secreted factors above the level of detection. To our surprise, we found no difference between the secretome profile of HB ACM compared to AB ACM for anti-inflammatory cytokines or pro-inflammatory cytokines (Fig. 3A, B); the AB ACM and the HB ACM were also similar to the ACM cytokine profile from an independent healthy control iPSC line (data not shown). In contrast, we found that HB MCM produced significantly higher levels of the anti-inflammatory factors IL-1RA and IL-10 compared to the AB MCM (Fig. 3C) as well as an unrelated control iPSC line (data not shown), but there was no difference between HB MCM and AB MCM in the key inflammatory molecule MCP-1 (Fig. 3D). Although FGF2 was significantly upregulated in the AB MCM (Fig. 3D), its overall expression level was low compared to MCP-1.

**Figure 3:**
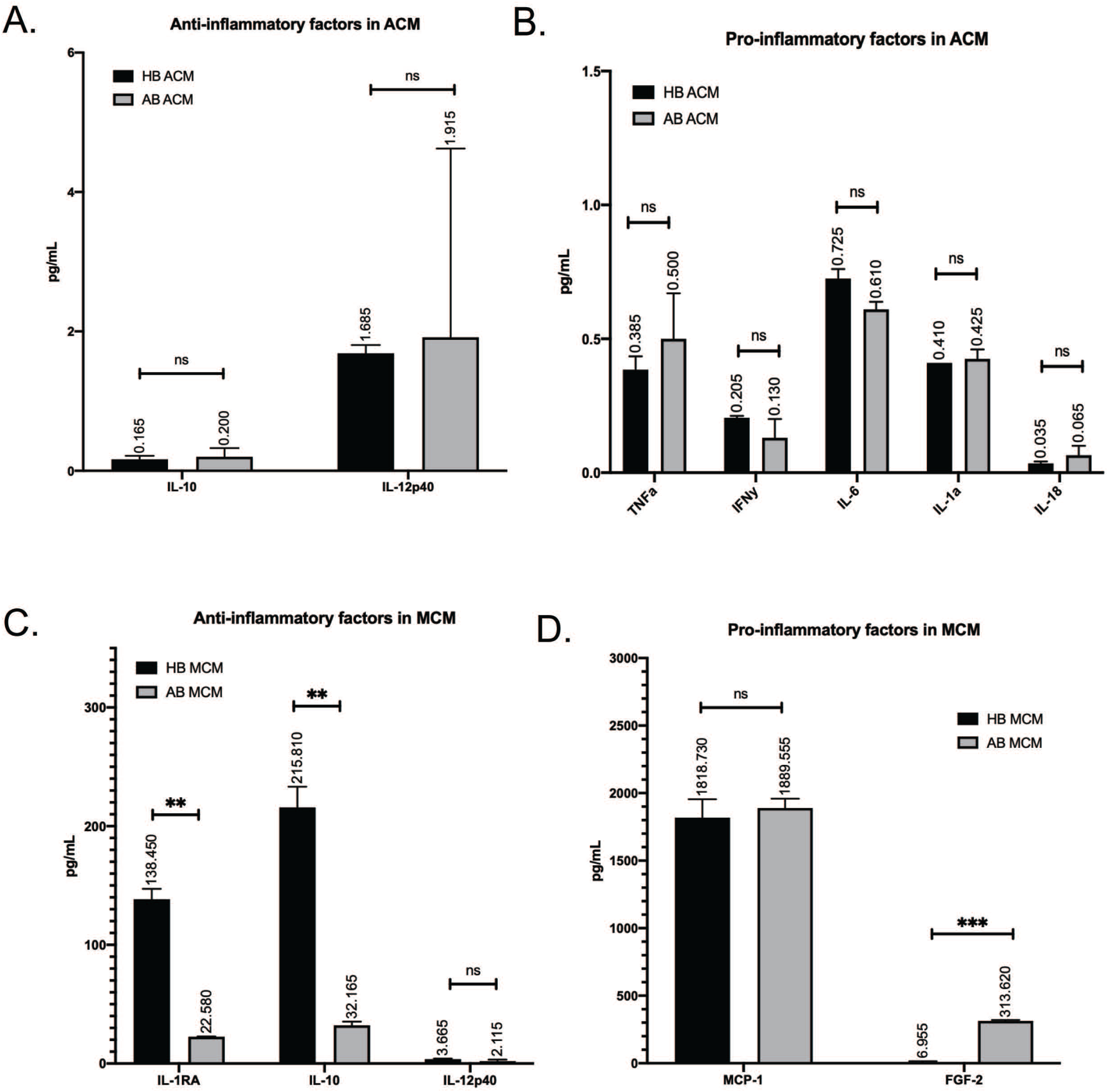
Differing secretome profiles identified between HB and AB microglia. (**A**) Multiplex cytokine array on HB (black bar) and AB (gray bar) ACM reveal similar levels of secreted anti-inflammatory cytokines IL-10 and IL12p40 [t-tests, ns=0.7517, 0.9155]. (B) Similar levels of astrocyte-relevant pro-inflammatory cytokines TNFa, IFNy, IL-6, IL-1a, and IL-18 were also noted between HB and AB ACM [t-tests, ns=0.4547, 0.2741, 0.0695, 9.6094, 0.3604]. (C) Multiplex cytokine array on HB (black bar) and AB (gray bar) MCM reveals significantly higher secreted levels of anti-inflammatory cytokines IL-1RA, IL-10, and a trend for higher IL12p40 by HB microglia compared to AB microglia [t-tests, **p=0.0028, **p=0.0046, ns=0.2348]. (D). Pro-inflammatory cytokines MCP-1, FGF-2, and IL12p70 are found to be secreted at higher levels in AB MCM than HB MCM [t-tests, ns= 0.5795, ***p=0.0006, ns=0.5631].

We next treated differentiated motor neurons with either HB ACM or AB ACM for 48hrs. Western blot analysis showed that neither HB ACM nor AB ACM altered NF protein levels for either HB or AB motor neurons (Fig. 4A-G). Interestingly, we did observe that both HB ACM and AB ACM induced a significant increase in phNF puncta in the HB motor neurons (Fig. 4H, I), whereas neither HB ACM nor AB ACM induced a change in phNF puncta in the AB motor neurons (Fig. 4J, K) again despite no difference in total NF200 expression for HB and AB motor neurons treated with either HB ACM or AB ACM (Fig. 4L, M).

**Figure 4:**
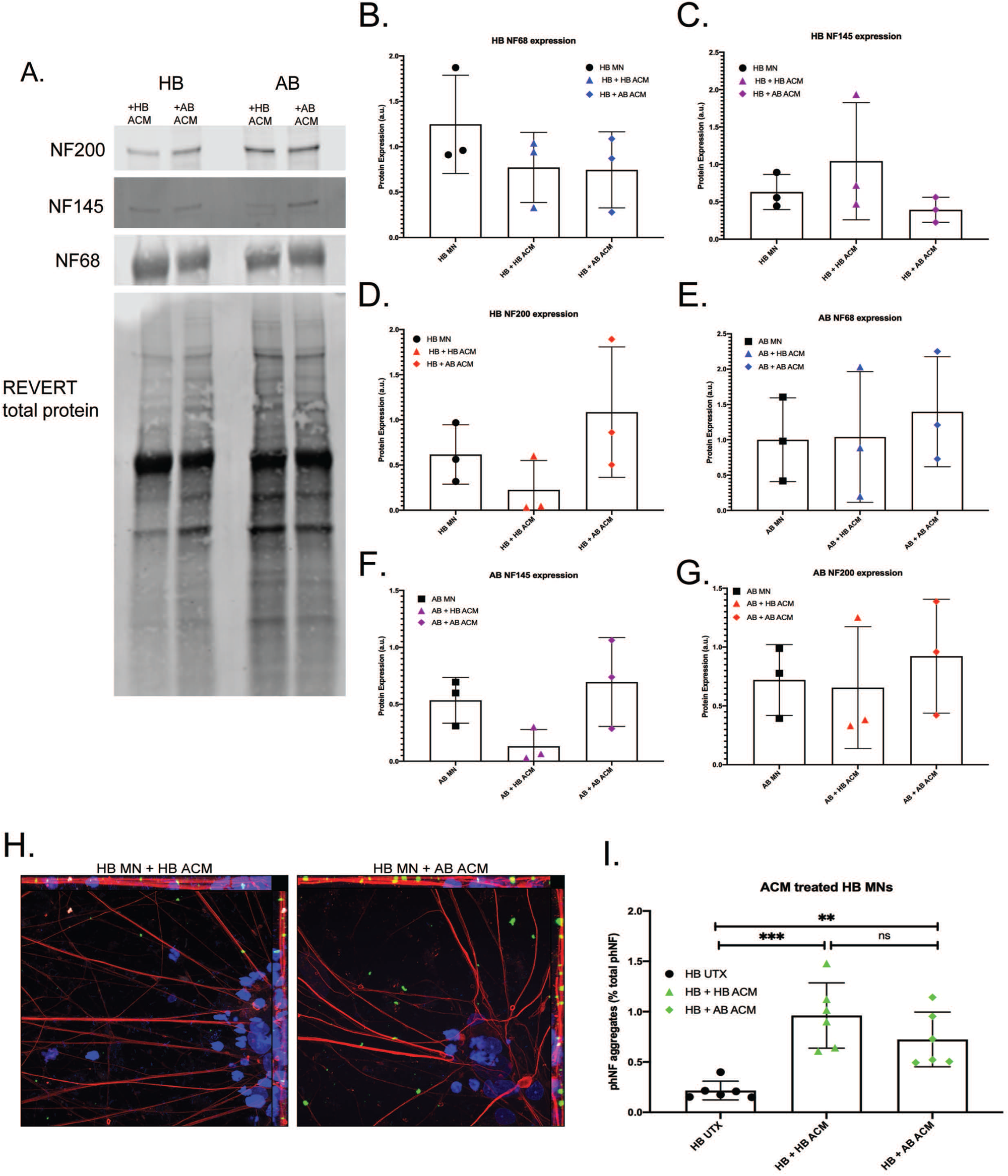
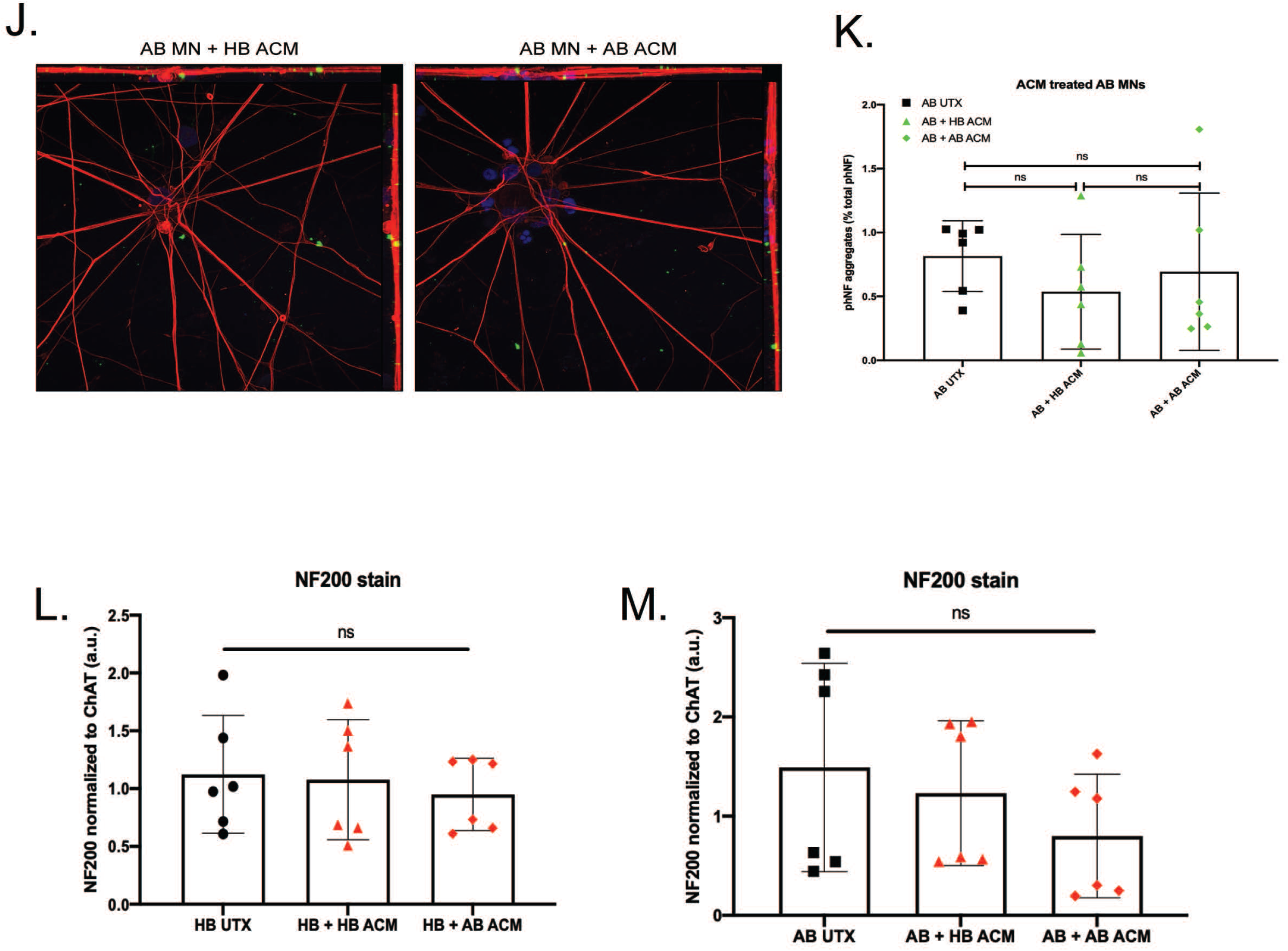
Effect of HB and AB astrocyte conditioned media on NF subunit expression and phNF+ aggregation. (**A-G**) WB for NF68, NF145, and NF200 subunits reveal no significant changes after treatment with HB or AB astrocyte conditioned media (ACM) on HB MNs (**B-D**) or AB MNs (**E-G**). Untreated (UTX, black) samples are copied from Fig. 2 for ease of comparison to ACM treatment. (**H**) Representative images of HB MN ICC after HB (left) or AB (right) ACM treatment (red = NF200, green = phNF, blue = DAPI, 63x objective). (**I**) Quantification for phNF in HB MNs shows significant increase in phNF+ aggregates after ACM treatment from either HB or AB ACM with no significant difference between ACM treatments [1-way ANOVA, ***p=0.0003, **p=0.0083, ns= 0.2579]. (**J**) Representative images of AB MNs after HB (left) or AB (right) ACM (red = NF200, green = phNF, blue = DAPI, 63x objective). (**K**) Quantification showed no change in phNF+ aggregation in AB MNs after treatment with either HB or AB ACM [1-way ANOVA, ns=0.5683, 0.8932, 0.8335]. ICC for NF200 in HB (**L**) and AB (**M**) MNs confirms no significant changes after HB ACM or AB ACM treatment.

We next tested how the addition of MCM would further impact NF expression and accumulation. IL-10 and IL-1 receptor antagonist (IL-1RA) prevent glial hyperactivation and IL-1 activation, respectively, thereby reducing overall inflammatory signaling [42, 43]. Therefore, because we saw a significant increase of IL-10 and IL-1RA production in the HB MCM and no difference between HB ACM and AB ACM (Fig. 3, 4), we tested whether the addition of HB MCM to AB ACM would decrease motor neuron expression of phNF aggregates. We pre-treated AB astrocytes with HB MCM for 48hrs. We then changed the medium, collected AB ACM 48hrs later, and added the HB MCM treated AB ACM to either HB or AB motor neurons. HB MCM-AB ACM treatment had no effect on overall NF200 expression in the HB motor neurons (Fig. 5A,B). Interestingly, this treatment did not show an increase in phNF puncta in the HB motor neurons (Fig. 5A, C), which is in contrast to what was observed with AB ACM treatment alone (Fig. 4I). HB MCM-AB ACM treatment on the AB motor neurons also did not induce a change in overall NF200 expression (Fig. 5D, E), but this treatment did significantly reduce the presence of phNF puncta in the AB motor neuron cultures (Fig. 5D, F). Together, these data suggest that factors released from the HB microglia are involved in modulating phNF deposition in ALS motor neurons.

**Figure 5:**
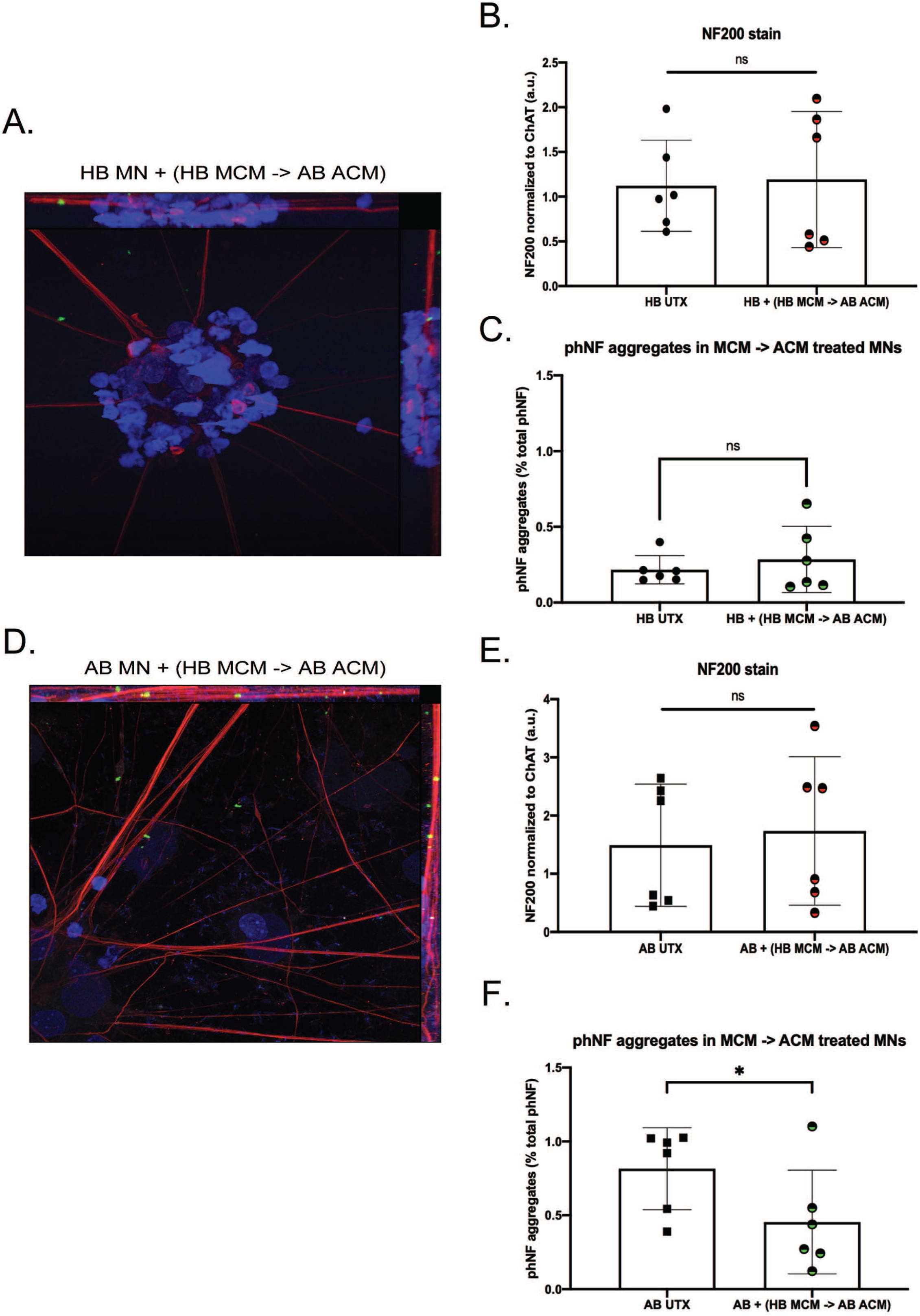
Effect of HB and AB microglia conditioned media on ACM induced ALS phenotype. (**A**) Representative image of HB motor neurons (MNs) treated with medium from AB astrocytes pre-treated with HB microglia conditioned media (HB MCM) (red= NF200, green= phNF, blue = DAPI 63x objective). (**B**) Quantification confirms no change in NF subunit expression in HB MNs following treatment with AB ACM pre-treated with HB MCM [t-test, ns=0.7795]. (**C**) Quantification shows prevention of ACM induced phNF+ aggregates in HB MNs after pre-treatment of AB astrocytes with HB MCM [t-test, ns=0.4340]. (**D**) Representative image of AB MNs treated with AB ACM pre-treated with HB MCM (red= NF200, green= phNF, blue = DAPI 63x objective) (**E**) Quantification confirms no change in NF subunit expression in AB MNs after application of AB ACM pre-treated with HB MCM [t-test, ns=0.3222]. (F) Quantification reveals a significant reduction of phNF+ aggregates when HB MCM pre-treated AB ACM is applied to AB MNs [t-test, *=0.0203].

## Discussion

ALS is an always fatal disease that currently lacks effective treatment options. Vast amounts of work have focused on elucidating the molecular mechanisms associated with motor neuron loss in order to identify novel therapeutic targets. Among these studies, data suggest that both phNF aggregation and glial-neuron interactions can contribute to motor neuron malfunction and loss in ALS [1, 23]. In the current study, we found that glial produced factors can augment phNF accumulation suggesting a link between these two pathogenic pathways.

Axonal swellings were identified in postmortem tissue from a motor neuron disease patient in the late 1960s [44] with subsequent studies identifying neurofilament accumulation as a pathological property of ALS [5]. However, there are contrasting data across in vitro, in vivo, and human studies regarding whether NF levels are upregulated or downregulated in ALS and the relationship to disease. For example, neuronal NF overexpression in mice induces an ALS-like phenotype [45, 46], NF deletion in SOD1 mice is protective [47], and NF levels are significantly increased in the CSF and blood of ALS patients [12]. In contrast, others have found that NF levels are decreased in mutant SOD1 iPSC-derived motor neurons [20] and crossing mutant SOD1 mice with mice overexpressing NF-L or NF-H reduces disease phenotypes and extends lifespan [48, 49]. Although we find significantly elevated transcript levels of NF-L, NF-M, and NF-H in the AB motor neurons (Fig. 1), this did not translate into elevated protein levels (Fig. 2). Nevertheless, we observed an increased abundance of phosphorylated NF accumulations in the AB motor neurons (Fig. 2). As such, it may be less about the overall protein abundance and more about altered subunit dynamics leading to aggregation and disrupted downstream signaling leading to hyperphosphorylation [11, 20]. As NFs are essential for structural stability, nerve conduction velocity, organelle interactions, and synaptic function [50], even subtle disruptions could have profound consequences for neuronal health and survival.

Astrocytes and microglia are integral components of a normally functioning nervous system. Astrocytes, the most abundant cell type in the central nervous system, perform a wide range of functions including providing trophic support to surrounding cells, maintaining extracellular ion balance, secreting molecules necessary for neuronal energy and metabolism, and taking up and recycling neurotransmitters [51]. Microglia, the resident immune cells of the central nervous system, continuously survey the environment to maintain synaptic function and debris clearance [52]. However, injury or disease can cause both astrocytes and microglia to alter their function and upregulate inflammatory signaling. In this regard, studies have found that both mutant SOD1 expressing mice and mutant OPTN expressing mice upregulate NFκB expression that drives production of IL-6, IL-1β, and TNFα from both astrocytes and microglia [31, 53], which overlaps with cytokines observed in the iPSC-derived ACM (Fig. 3). Previous studies have clearly demonstrated that familial and sporadic ALS iPSC-derived astrocytes induce motor neuron loss [54], but much less is known about the specific astrocyte secreted proteins involved, the toxic nature of the ALS microglial secretome, or how combined astrocyte-microglial signaling influence motor neuron health and survival in ALS.

What we found surprising, though, was that the astrocyte cytokine profile for the HB was very similar to the AB (Fig. 3). Given the robust toxic effects of ALS astrocytes in other models, we hypothesized that we would find an increase in pro-inflammatory cytokines and/or a decrease in the anti-inflammatory cytokines in the AB ACM compared to the HB ACM. However, since we did not see a difference between the ACM cytokine profiles, it was then not surprising that both the HB ACM and the AB ACM induced an upregulation of phNF in the HB motor neurons (Fig. 4). These data suggest that the HB motor neurons are capable of developing an ALS-like phenotype, but perhaps other systemic signaling processes are keeping them from tipping to a more pathological state. This idea is supported by our data showing that pre-treating astrocytes with HB MCM reduced the phNF deposition (Fig. 5). Although the cytokine data presented here (Fig. 3) focused on a small number of factors that were above the level of detection in the assay, it is likely that there are other factors involved in the microglia-to-astrocyte signaling. In this regard, additional analysis of the motor neuron RNAseq data (Fig. 1) as well as generating RNAseq data from ACM-MCM treated motor neurons will be informative for future pathway analysis studies.

Finally, our study focused on a unique discordant identical twin pair to model sporadic ALS using the HB as an approximation of an isogenic cell line. Therefore, we are not able to generalize these data to other sporadic ALS cases or to familial ALS cases. Future studies will be needed to better characterize the glial-motor neuron interplay in ALS more broadly, but our data provide an interesting starting point for assessing how microglial-astrocyte-motor neuron interactions contribute to ALS pathology.

## Acknowledgements

This work was funded in part by the Phoebe Lewis Regenerative Medicine Fund and grants from the Neuroscience Research Center and the Center for Immunology at the Medical College of Wisconsin. We thank the Froedtert Hospital ALS Clinic for initially providing the patient samples and Dr. Mike Dwinell for helpful discussions.

## Author contributions

Conceptualization, RLA, ADE; Methodology, RLA, JWA, JA, RAU, MTZ, AJM; Data Analysis, RLA, JWA, JA, MTZ, AJM; Resources, RAU, ADE; Writing – Original Draft Preparation, RLA, JWA, ADE; Writing – Review & Editing, RLA, JWA, JA, RAU, MTZ, AJM, ADE; Funding Acquisition, ADE

## Figure Legends

**Figure S1:**
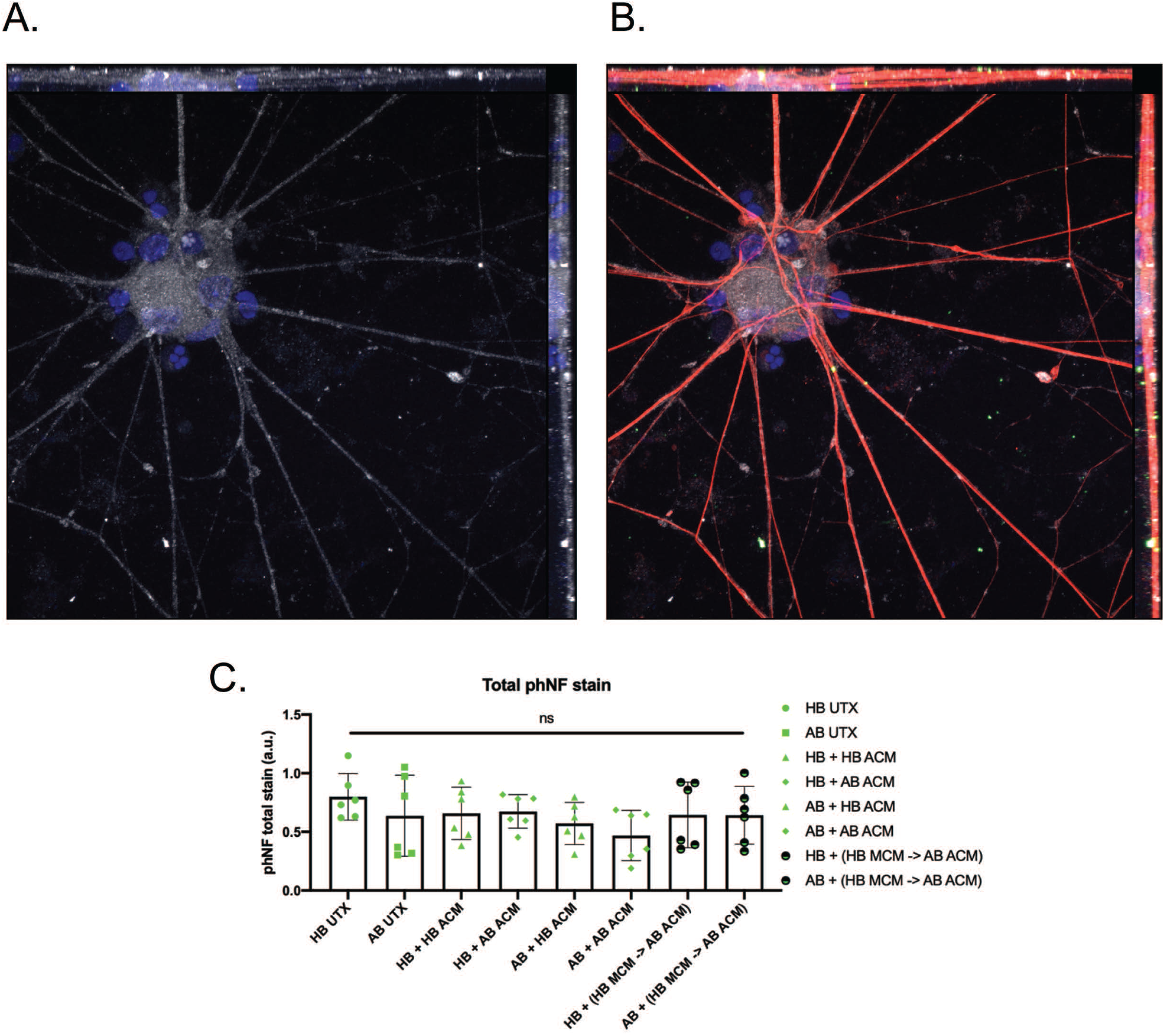
Only phNF+ aggregates differ between treatment groups –not total phNF– indicative of an ALS phenotype. (A) Representative image of HB MNs to show that MNs quantified in ICC analyses were ChAT+ (blue = DAPI, white = ChAT, 63x objective). (B) NF200 and phNF+ aggregates colocalized with ChAT staining (blue = DAPI, white = ChAT, red = NF200, green = phNF, 63x objective). (C) Quantified ICC of total phNF shows no differences between HB and AB MNs, and no significant changes between treatment groups [1-way ANOVA, ns].

**Figure S2:**
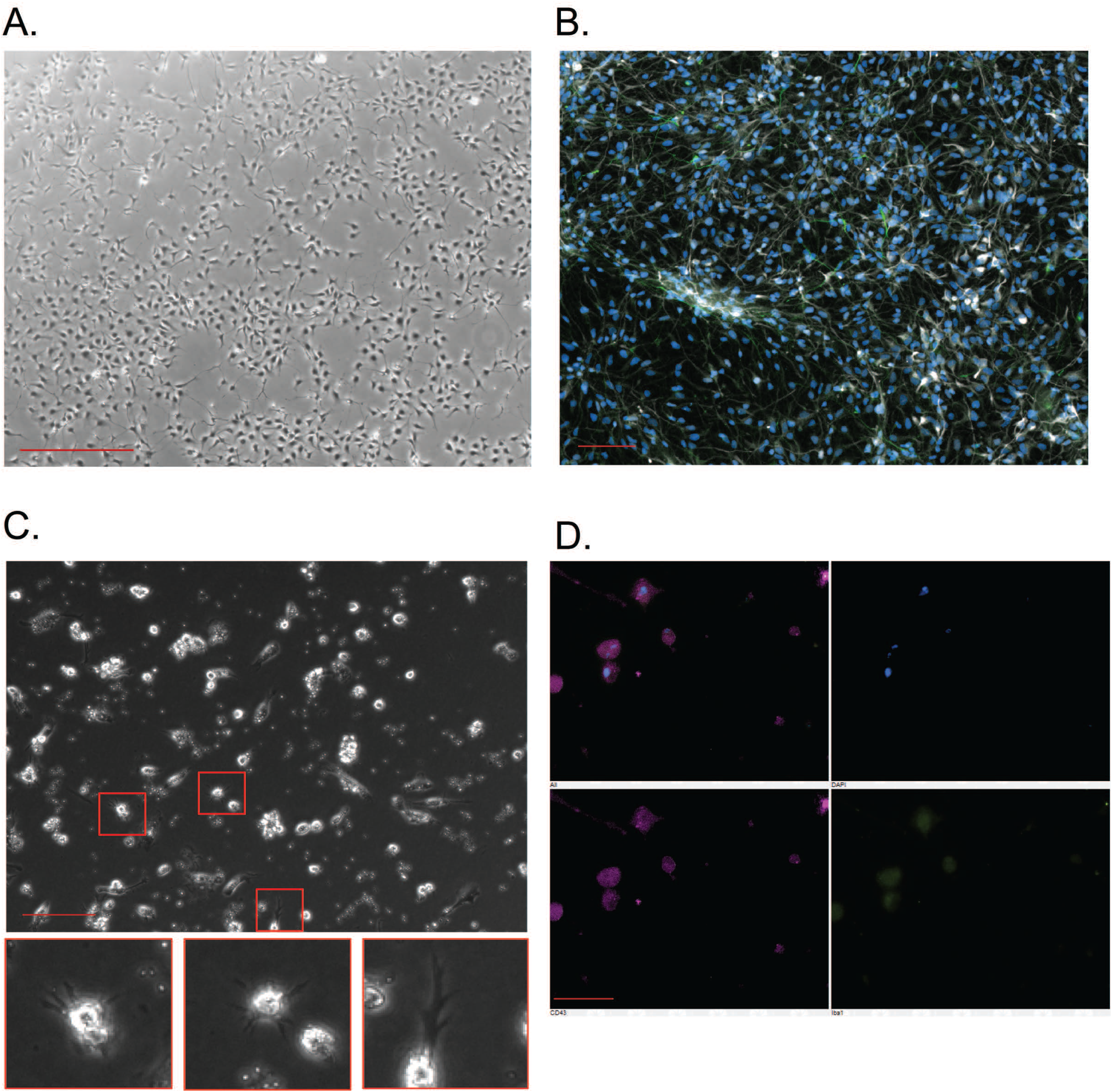
Representative images of astrocytes and microglia differentiated from iPSCs. (**A**) Brightfield image of AB astrocytes at p.3 (4x objective; scale bar = 500µm). (**B**) ICC for canonical astrocyte markers GFAP (green) and S100B (white) in AB astrocytes after p.4 (10x objective, blue = DAPI, scale bar = 100µm). (**C**) Brightfield image of iPSC-derived microglia on day 30 of differentiation (20x objective, scale bar = 50µm). Red boxes outline individual microglia, shown in the zoomed images below. (**D**) ICC for canonical microglia markers CD43 (purple, bottom left) and Iba1 (green, bottom right) in iPSC-derived microglia (60x objective, scale bar = 10µm) Top left panel: merged image, top right: blue = DAPI.

